# Hippo Signaling Pathway Activation during SARS-CoV-2 Infection Contributes to Host Antiviral Response

**DOI:** 10.1101/2022.04.07.487520

**Authors:** Gustavo Garcia, Yijie Wang, Joseph Ignatius Irudayam, Arjit Vijey Jeyachandran, Sebastian Castillo Cario, Chandani Sen, Shen Li, Yunfeng Li, Ashok Kumar, Karin Nielsen-Saines, Samuel W. French, Priya S Shah, Kouki Morizono, Brigitte Gomperts, Arjun Deb, Arunachalam Ramaiah, Vaithilingaraja Arumugaswami

**Affiliations:** Department of Molecular and Medical Pharmacology, University of California, Los Angeles, CA 90095, USA; Division of Cardiology, Department of Medicine, David Geffen School of Medicine, UCLA, Los Angeles, CA 90095, USA; UCLA Children’s Discovery and Innovation Institute, Mattel Children’s Hospital UCLA, Department of Pediatrics, David Geffen School of Medicine, UCLA, Los Angeles, CA 90095, USA; Translational Pathology Core Laboratory, Department of Pathology and Laboratory Medicine, David Geffen School of Medicine, UCLA, Los Angeles, CA 90095, USA; Department of Ophthalmology, Visual and Anatomical Sciences, Wayne State University, Detroit, MI USA; Jonsson Comprehensive Cancer Center, UCLA, Los Angeles, CA 90095, USA; Department of Chemical Engineering, University of California, Davis, CA 95616, USA; Division of Hematology and Oncology, Department of Medicine, David Geffen School of Medicine, University of California, Los Angeles, CA 90095, USA; UCLA AIDS Institute, David Geffen School of Medicine, University of California, Los Angeles, CA 90095, USA; Eli & Edythe Broad Center of Regenerative Medicine and Stem Cell Research, UCLA, Los Angeles, CA 90095, USA; Molecular Biology Institute, UCLA, Los Angeles, CA 90095, USA; California Nanosystems Institute, UCLA, Los Angeles, CA 90095, USA; Tata Institute for Genetics and Society, Centre at inStem, Bangalore, KA 560065, India; Department of Ecology and Evolutionary Biology, University of California, Irvine, CA 92697, USA; Section of Cell and Developmental Biology, University of California, San Diego, CA 92093, USA

**Keywords:** SARS-CoV-2, COVID-19, Hippo signaling pathway, Antiviral, YAP/TAZ, LATS1/2, MST1/2

## Abstract

SARS-CoV-2, responsible for the COVID-19 pandemic, causes respiratory failure and damage to multiple organ systems. The emergence of viral variants poses a risk of vaccine failures and prolongation of the pandemic. However, our understanding of the molecular basis of SARS-CoV-2 infection and subsequent COVID-19 pathophysiology is limited. In this study, we have uncovered a critical role for the evolutionarily conserved Hippo signaling pathway in COVID-19 pathogenesis. Given the complexity of COVID-19 associated cell injury and immunopathogenesis processes, we investigated Hippo pathway dynamics in SARS-CoV-2 infection by utilizing COVID-19 lung samples, and human cell models based on pluripotent stem cell-derived cardiomyocytes (PSC-CMs) and human primary lung air-liquid interface (ALI) cultures. SARS-CoV-2 infection caused activation of the Hippo signaling pathway in COVID-19 lung and *in vitro* cultures. Both parental and Delta variant of concern (VOC) strains induced Hippo pathway. The chemical inhibition and gene knockdown of upstream kinases MST1/2 and LATS1 resulted in significantly enhanced SARS-CoV-2 replication, indicating antiviral roles. Verteporfin a pharmacological inhibitor of the Hippo pathway downstream transactivator, YAP, significantly reduced virus replication. These results delineate a direct antiviral role for Hippo signaling in SARS-CoV-2 infection and the potential for this pathway to be pharmacologically targeted to treat COVID-19.

## INTRODUCTION

Severe acute respiratory syndrome-related coronavirus 2 (SARS-CoV-2) is a betacoronavirus of zoonotic origin having similarity with bat SARS-CoV-like viruses. SARS-CoV-2 is responsible for the outbreak of the Coronavirus Disease 2019 (COVID-19) pandemic leading to over 5 million deaths and 270 million cases worldwide^1–3^. SARS-CoV-2 primarily causes severe lung injury with diffuse alveolar damage resulting in the development of acute respiratory distress syndrome (ARDS)^4^. The virus is also known to infect heart, kidney, brain and digestive systems due to the widespread expression of host cell surface membrane receptor angiotensin converting enzyme 2 (ACE2) and other receptors, including NRP1, AXL and CD147^5–11^. Therefore, patients develop additional systemic complications including acute kidney injury^12–14^, vascular inflammation (endotheliitis), and cardiac effects^15,16^. Viral replication-mediated epithelial cell injuries result in a vigorous immune response leading to organ failure, and possibly death in severe cases. Patients with various underlying conditions such as cardiovascular diseases, diabetes, and obesity are linked with increased risk of COVID-19 mortality^17,18^.

To replicate, viruses hijack cellular machinery and signaling pathways. Thus, understanding the molecular basis of dysregulated signaling circuitry can provide potential therapeutic targets. Utilizing a library of kinase inhibitors, we previously demonstrated critical pathways that are important for SARS-CoV-2 virus replication such as the DNA-Damage Response (DDR), mTOR-PI3K-AKT, and ABL-BCR/MAPK pathways^19^. COVID-19 pathogenesis is a complex process involving crosstalk with multiple cellular pathways. To understand the fundamental principles underlying RNA viral mediated cell injury processes, we hypothesize that the conserved cellular circuitry of the Hippo signaling pathway, which is involved in tissue growth, immune response, and inflammation, can contribute to regulating SARS-CoV-2 replication and COVID-19 pathogenesis. Previously, we observed that the Zika Virus (ZIKV), a member of the *Flaviviridae* family, dysregulates this pathway^20^. The core components of the Hippo pathway in mammals are STE20-like kinases STK4/MST1 and STK3/MST2 (Hpo in *Drosophila*), large tumor suppressors (LATS1/2, Wts in *Drosophila*), MOB kinase activator 1A and 1B (Mats in *Drosophila*), Neurofibromatosis 2 (NF2), and Salvador 1 (SAV1)^21^. MST1/2 and LATS1/2 are serine/threonine kinases, whereas NF2 and SAV1 are adaptors or co-factors. Upon activation by stimuli, MST1/2 phosphorylate and activates the downstream serine/threonine kinase LATS1/2. LATS1/2 can be phosphorylated by MAP4Ks independent of MST1/2 axis. The phospho-LATS1/2, along with SAV1, then phosphorylates YAP (Yes-associated protein), a transcription co-activator and oncogene. Phospho-YAP, bound to a 14-3-3 protein and retained in the cytoplasm, is subsequently ubiquitinated and degraded by proteasomes. While the Hippo pathway is inactive, the unphosphorylated YAP and its homologous partner TAZ translocate to the nucleus, where they bind to the TEAD family of transcription factors and mediate expression of target genes involved in cell survival and proliferation (Birc5, Id, CTCF, Axin2, Myc, CycD, etc.)^22^. As the Hippo signaling pathway controls progenitor cell proliferation and differentiation, and the size of organs^23–27^, the pathway is closely linked to mitochondrial energy metabolism and biogenesis. Studies suggest that the Hippo pathway modulates host antiviral immune responses^28–33^, where YAP inhibits antiviral defense mechanisms by antagonizing the function of pro-innate immune factor TBK1^34^. Inherited autosomal recessive mutations of MST1 (STK4) in human have been attributed to primary immunodeficiency with T- and B-cell lymphopenia, neutropenia, and defective regulatory T cells^30,35–37^. Thus, we postulate that Hippo signaling pathway plays a critical role in SARS-CoV-2 antiviral response and COVID-19 pathogenesis.

## RESULTS/DISCUSSION

To understand the pathophysiological effect of SARS-CoV-2 on the Hippo signaling pathway, first we performed transcriptomic analysis of RNA sequencing (RNA-seq) data sets of lung samples from COVID-19 patients (n=5). KEGG pathway analysis indicates that many genes (45 genes) in the Hippo pathway were differentially regulated (*p* <0.01) (Figure 1A), and 33 (73%) of these genes are upregulated, likely due to an antiviral host response. WWTR1/TAZ, a YAP homologue and downstream transcriptional coactivator of the Hippo signaling pathway, was found to be significantly downregulated. Phosphorylation of YAP (phospho-YAP) at Serine (Ser or S) 127 is a hallmark of Hippo pathway activation and results in cytoplasmic retention and inactivation of its transactivation function. We investigated YAP/TAZ phosphorylation status by immunohistochemical staining and analysis of autopsy lung tissue samples from COVID-19 patients and control lungs. Using RNA-fluorescence *in situ* hybridization (RNA-FISH), SARS-CoV-2 viral genomes were detected in the COVID-19 lung tissue (Figure 1B). In addition, immunohistochemistry demonstrated a higher level of phospho-YAP (Ser127) in the COVID-19 lung compared to control lung (Figure 1B). The COVID-19 patient lung tissue also had a large amount of inflammatory cellular infiltration. (Supplementary Figure 1A).

**Figure 1.**
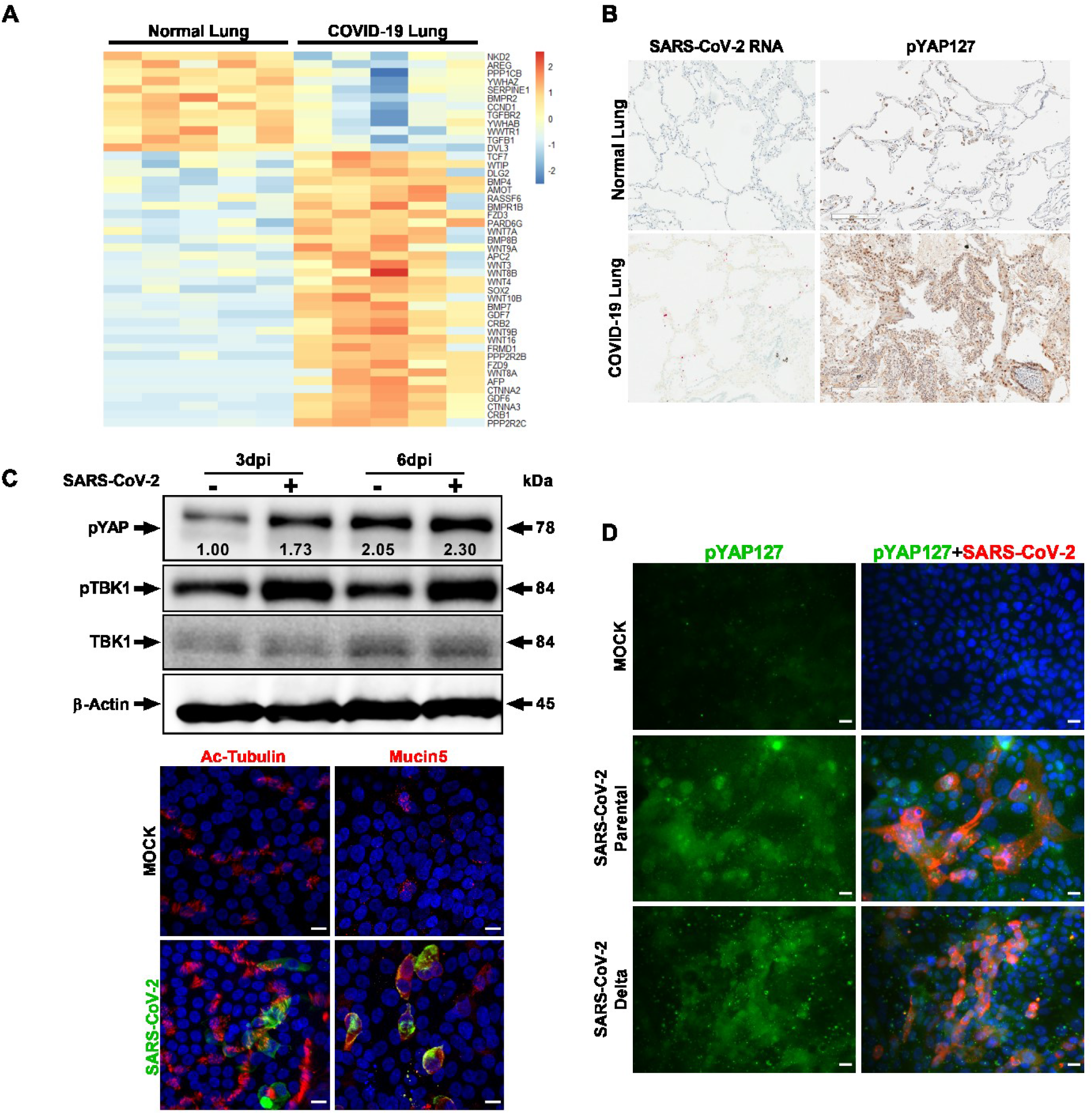
Hippo signaling pathway is activated in lung tissues of COVID-19 patients and infected human lung *in vitro* cell cultures. (A) Transcriptome analysis of five negative control and autopsy lung samples from patients deceased due to SARS-CoV-2 infection are depicted. KEGG pathway database was used to examine the Hippo signaling pathway genes in the differentially expressed genes (DEGs). Heatmap shows the expression levels of the 45 DEGs (*p* <0.01) involved in Hippo signaling pathway. Blue and red colors represent downregulated and upregulated genes, respectively. (B) Immunohistochemistry (IHC) of COVID-19 lung autopsy tissue shows high level of phospho-YAP Ser127 protein (dark brown). SARS-CoV-2 RNA (red) presence in COVID-19 lung was confirmed by RNA-FISH (RNAscope). Images are obtained at 20x magnification. (C) Western blot analyses of lung ALI cells show activation of Hippo-TBK1 pathways during SARS-CoV-2 infection. Proximal lung air-interface cultures are susceptible to SARS-CoV-2 infection. Antibody targeting Spike protein was used for probing infection by immunohistochemistry. Cell-specific markers, such as Ac-tubulin (ciliated cells) and Mucin5 (mucus cells) were detected by antibody probes to define infected cell types. Scale bar 10 μm. (D) Human airway epithelial cells (Calu-3) were analyzed with IHC for pYAP127 and SARS-CoV-2 Spike protein in uninfected (Mock) and infected cells. Parental and Delta strains were used for infection studies. Scale bar: 25 μm. Representative data from three independent studies are provided.

In order to further investigate SARS-CoV-2 viral replication and the Hippo signaling pathway, we utilized a biologically relevant primary human proximal airway cell culture system, which consisted of mucociliary epithelial cells grown at an air-liquid interface (ALI). Human lung airway basal stem cells (ABSC) harvested from the trachea and bronchi of healthy lungs were differentiated into a ciliated pseudostratified columnar epithelium in a collagen-coated transwell membrane culture system^19,38,39^. The ciliated and secretory airway epithelial cell types in the ALI culture were verified using immunostaining for the cell type-specific markers, acetylated tubulin and Mucin5AC, respectively (Figure 1C). The apical surface of the ALI cultures was exposed to SARS-CoV-2 infection to understand how the virus affects the Hippo pathway at a post-translational level. Parental SARS-CoV-2 (Isolate USA-WA1/2020 at MOI of 0.01) from BEI Resources was used for infection studies in the UCLA BSL3 high-containment facility. Western blot analysis of protein samples collected at 3 and 6 days post-infection (dpi), showed an increased level of phosphorylated YAP (Ser127) suggesting activation of the Hippo signaling pathway during SARS-CoV-2 infection (Figure 1C). In addition, we observed a concurrent activation of the innate immune pathway as evidenced by an increase in phosphorylated TBK1 (Ser172). SARS-CoV-2 infection was confirmed in the lung ALI culture by immunohistochemistry, which revealed both ciliated and mucus secreted cells were infected at 6 dpi (Figure 1C). Next, we examined the Hippo signaling pathway in the Calu-3 human airway epithelial cell line during infection with SARS-CoV-2 parental as well as the Delta Variant of Concern (VOC) (Figure 1D). We observed that both the parental and Delta variant strains of SARS-CoV-2 increased phospho-YAP (Ser127) level in the infected Calu-3 cell cultures. Interestingly, the phospho-YAP (Ser127) protein aggregates formed punctate structures in the cytoplasm, likely for potential degradation in autophagosomes^40^. Taken together, the Hippo signaling pathway is activated in COVID-19 infected lungs and SARS-CoV-2 infected *in vitro* lung culture systems.

Although SARS-CoV-2 is a major respiratory pathogen, COVID-19 manifestations in the cardiovascular system are also well documented^17,18,41–43^. To further understand how the Hippo signaling pathway may be modulated by SARS-CoV-2 and the host response, we studied the pathway in cardiomyocytes. Human pluripotent stem cell-derived cardiomyocytes (hPSC-CMs) have been shown to be efficient at recapitulating cardiovascular diseases at a cellular level^44,45^, have demonstrated susceptibility to SARS-CoV-2 replication^46^, and have been used as a platform to study potential therapeutics against SARS-CoV-2 infection^19^. Transcriptomic analysis of the SARS-CoV-2 infected hPSC-CM system showed that many genes in the Hippo pathway are significantly dysregulated compared to uninfected mock hPSC-CM control (*p* <0.01) (Supplementary Figure 1B). While comparing the Hippo pathway DEGs of COVID-19 lung samples (Figure 1A) and of iPSC-CMs (Supplementary Figure 1B), we observed that genes such as BMP7, PPP2R2B, CTNNA3, GDF6 and SERPINE1 are commonly deregulated. Therefore, we used the hPSC-CM to further investigate the kinetics of the Hippo signaling pathway during viral infection. SARS-CoV-2 infected cardiomyocytes were confirmed by specific cell-staining of cardiac troponin T (cTnT) and viral Spike proteins (Figure 2A). In addition, SARS-CoV-2 induced cell injury and apoptosis by activation of cleavage of caspase-3 (CC3) indicated the high susceptibility of hPSC-CM to cell death by infection. Western blot confirmed that YAP1 protein is degraded at 48 hours post-infection (hpi) (Figure 2B). Simultaneously, there is an induction of phospho-YAP1 (S127) at 2 hpi and 24 hpi. These two events indicate a strong activation of the Hippo signaling Pathway. Also, at 2 hpi, SARS-CoV-2 infection results in activation of the STAT1 Type I interferon (IFN) pathway (Figure 2B). Furthermore, immunohistochemical analysis showed that YAP/TAZ has a cytoplasmic localization in the infected cells (Figure 2C) and the level of pYAP127 has increased (Figure 2D). Phosphorylation at the Serine 127 residue leads to cytoplasmic distribution of the YAP protein and prevents its transactivation function. Collectively, our data conclude that the Hippo signaling cascade is also active during SARS-CoV-2 infection of cardiomyocyte systems, therefore we next sought to determine the functional relevance of this pathway.

**Figure 2.**
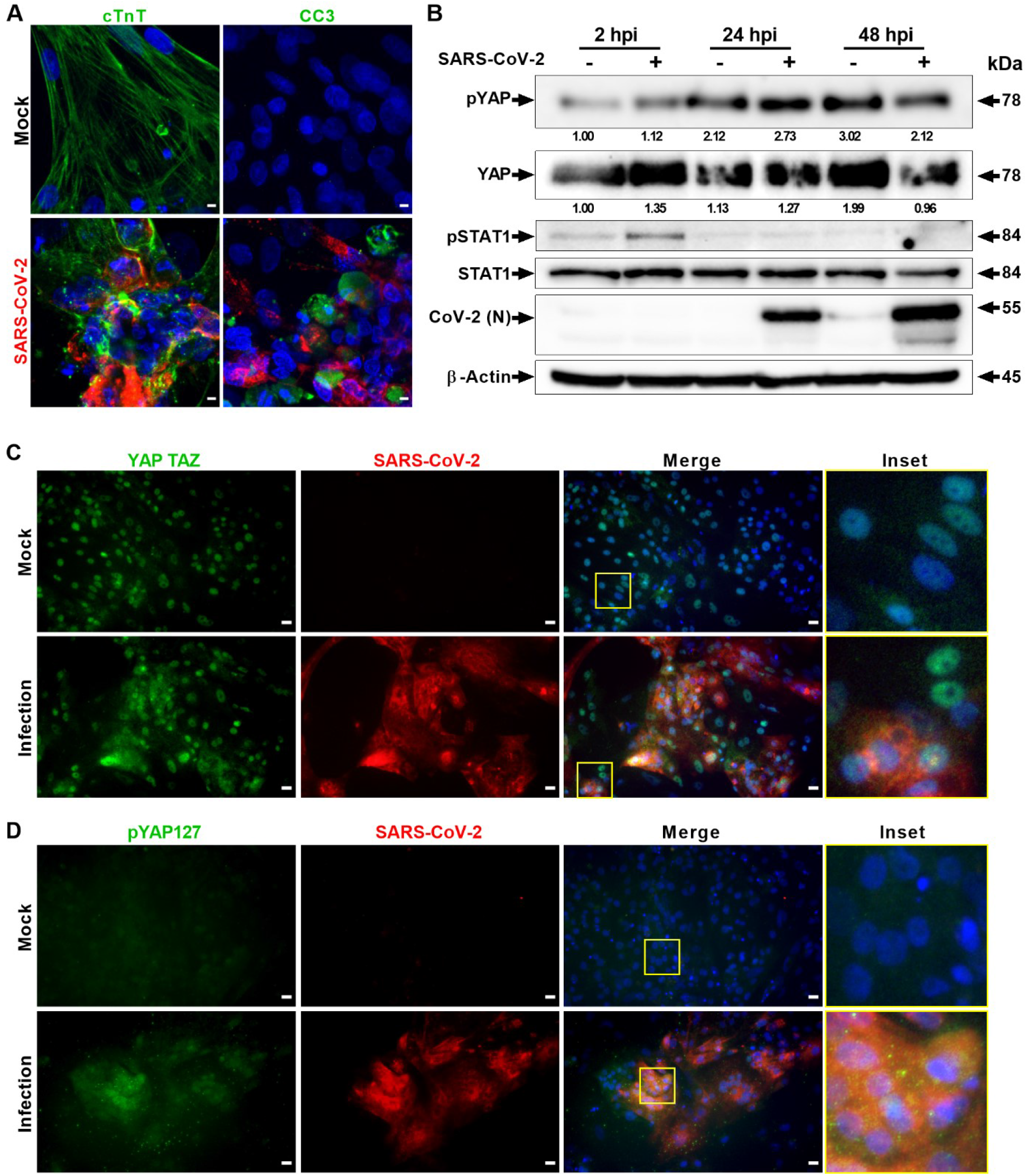
SARS-CoV-2 infection activates Hippo and antiviral STAT pathways in hPSC-CMs. (A) Confocal image analysis of SARS-CoV-2 (red) infected cardiomyocytes shows extensive damage to cTNT positive (green) cells, which undergo apoptotic cell death (green; cleaved caspase 3). Scale bar 5 μm. n=6 independent experiments. (B) Western blot analyzes show activation of Hippo and STAT1 pathways. Phospho-YAP127 level is increased at 2 and 24 hpi upon SARS-CoV-2 infection. N=2 independent experiments. (C) Immunohistochemistry analysis of SARS-CoV-2 infected PSC-CMs at 24 hpi reveals cytoplasmic localization of YAP/TAZ and (D) increase in pYAP127 level. Scale bar 25 μm.

In order to uncover the functional significance of the Hippo pathway, we utilized both genetic and pharmacological ablation approaches. The core Hippo signaling pathway is a kinase cascade where MST1/2 kinases and Sav1 form a complex to phosphorylate and activate LATS1/2 kinases. After activation, LATS1/2 can phosphorylate and inhibit YAP and TAZ transcription co-activators. YAP/TAZ, when dephosphorylated, interacts with TEAD1-4 and additional transcription factors to stimulate gene expression for cell proliferation and to inhibit apoptosis. Therefore, the Hippo pathway is regulated at multiple levels. MST1/2 and LATS1/2 are regulated upstream by other molecules like Merlin and KIBRA, and YAP/TAZ is regulated downstream by protein ubiquitination. Thus, we postulated that inhibition of MST1/2 and LATS1/2 would lead to an increase in YAP/TAZ activity, and consequently an increase in SARS-CoV-2 replication. To test this mechanism, we performed shRNA-mediated gene silencing experiments in hPSC-CM. For knockdown experiments, we transduced the cells with lentiviral vectors expressing non-specific control shRNA or shRNAs targeting YAP/TAZ or LATS1. 72 hours post-transduction, the cells were infected with SARS-CoV-2 at MOI of 0.01. At 48 hpi, the cells were fixed for IHC or harvested in RIPA protein lysis buffer for Western blot analysis. Immunohistochemical analysis revealed that the shRNA-mediated partial knock down of YAP1 gene in hPSC-CM cells resulted in significantly reduced SARS-CoV-2 infection, whereas LATS1 knockdown, increased SARS-CoV-2 infection (Figure 3A and D). The quantification of spike antigen positive infected cells is provided in Figure 3D. These observations were also verified at the protein level (Figure 3B). Taken together, our data provide evidence that the YAP/TAZ is a pro-viral factor, whereas LATS1 has antiviral function.

**Figure 3.**
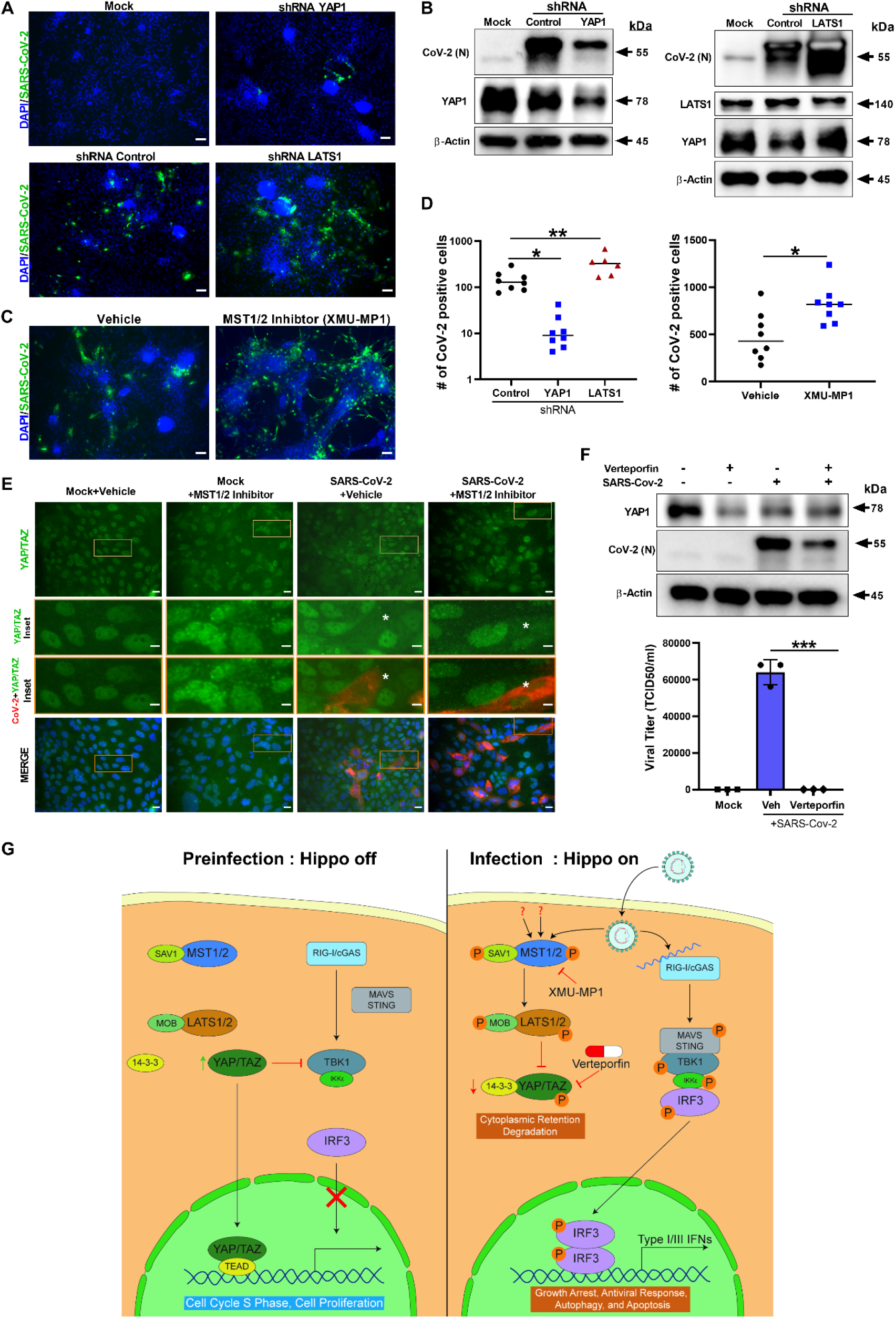
ShRNA-mediated knockdown and pharmacological modulation of SARS-CoV-2 replication. (A) IHC analysis of shRNA-mediated knockdown of YAP1 and LATS1 specific shRNAs showed efficiently reduced or increased SARS-CoV-2 replication (green) relative to shRNA control, respectively in hPSC-CMs. Scale bar 50 μm. (B) Western blot analysis of shRNA-mediated knockdown of YAP1 and LATS1 respective protein expression. (Con: Control shRNA). (C) IHC images of XMU-MP-1 (MST1/2 inhibitor) and vehicle treated hPSC-CMs are shown. Note: XMU-MP-1 increased SARS-CoV-2 replication (green) in hPSC-CM. (D) Graphs depict quantification of SARS-CoV-2 positive cells in infected hPSC-CM respective to panels A and C. Student T-test. **P >0.001. n=2 independent experiments. (E) IHC Images show YAP/TAZ protein (green) and SARS-CoV-2 Spike (red) in Calu-3cells. Note, MST1/2 inhibitor treated Calu-3 cells have higher number of infected cells. Inset and white asterisk hovers infected cells showing depletion of YAP/TAZ. Scale bar: 25 μm. Inset scale bar 10 μm. (F) Western blot analysis of Calu-3 cells treated with Verteporfin (1μM) and SARS-CoV-2 infection. Drug treatment resulted in reduction in SARS-CoV-2 infection. Graph shows the viral titer (TCID50/ml) measurement of infected as well as treated Calu-3 culture supernatant(representative data from two independent experiments) (G) Schematic diagram of our hypothetical model integrating Hippo and TBK1 signaling pathways during preinfection (Hippo off) and SARS-CoV-2 infection states (Hippo on). c-GAS, cyclic GMP-AMP synthase; IKKe, inhibitor of nuclear factor kappa B kinase subunit epsilon; MAVS, mitochondrial antiviral-signaling protein; RIG-I, retinoic acid inducible gene I protein; STING, stimulator of interferon response cGAMP interactor 1; TEAD, TEA domain transcription factors.

In addition, we investigated well characterized direct acting chemical inhibitors on the Hippo signaling pathway. XMU-MP-1 blocks MST1/2 kinase activities and thereby enhances YAP/TAZ activity. MST1 and MST2 are core components of the Hippo pathway and critical targets for tissue repair and regeneration^47^. Therefore, XMU-MP-1, a selective and reversible inhibitor, increases tissue repair and regeneration properties following tissue injuries^47,48^. XMU-MP-1 increases cardiomyocyte survival and reduces apoptosis following oxidative stress^48^. In mice, XMU-MP-1 has been shown to increase intestinal repair, liver repair and regeneration^47^, and preserve cardiac function and inhibit cardiomyocyte apoptosis in mice with transverse aortic constriction (TAC)^48^. Infection of cardiomyocytes and Calu-3 lung cells pre-treated with the MST1/2 inhibitor, XMU-MP-1 (10 μM), led to an increase in SARS-CoV-2 replication (Figure 3C, D and E). XMU-MP-1 compound is non-toxic to the Calu-3 cells (Supplemental Figure 2A). Interestingly, XMU-MP-1 treatment induced a coarse granular morphology to the nuclear localized YAP/TAZ (Figure 3E), and in infected cells the YAP/TAZ protein was depleted. Furthermore, XMU-MP1 treatment stimulated an increase in SARS-CoV-2 replication. Similar to the activity of XMU-MP-1, MST1/2 loss of function mutations in humans result in immune deficiency disorders and the affected individuals are more susceptible to human papilloma infections^33^. Consistent with the clinical observation, our study indicates that pharmacological inhibition of MST1/2 promotes the susceptibility of cardiac and lung cells to SARS-CoV-2 infection. Thus, MST1/2 or LATS1/2 loss of function mutations can potentially increase susceptibility to SARS-CoV-2 infections in humans.

Since YAP/TAZ is a proviral factor, pharmacological inhibition can provide additional therapeutic options for COVID-19 treatments. Thus, we focused on pharmacological modulation of YAP/TAZ. Verteporfin, a small molecule derivative of porphyrin, is a YAP-TEAD inhibitor. Verteporfin is a US Food and Drug Administration (FDA) approved drug used in patients with predominantly classic, subfoveal choroidal neovascularization (CNV) caused by AMD Age-related Macular Degeneration (VAM)^49^. In addition, because YAP is a pro-tumorigenic factor expressed in multiple cancers, Verteporfin has been used to inhibit YAP as an anti-cancer chemotherapeutic and adjuvant drug^50^, which increased phosphorylation of YAP1 (S127) in Calu-3 cells (Supplemental Figure 2B). Therefore, modulating YAP/TAZ with Verteporfin can provide more insight into YAP’s proviral properties. Calu-3 cells were pretreated with 1 μM Verteporfin (non-toxic concentration; Supplemental Figure 2A) for 24 hours and subsequently the cells were infected with SARS-CoV-2. At 48 hpi, the cells were harvested for protein analysis and the culture supernatants were collected for measuring viral titer. Verteporfin treatment resulted in a reduction of YAP/TAZ protein level compared to vehicle treated cells (Figure 3F) as well as a decrease in SARS-CoV-2 replication at 48 hpi. Viral titer (TCID50/ml) measurement of Verteporfin treated infected cell culture media revealed there was a significant decrease of viral production in Calu-3 cells (Figure 3F). Overall our results indicate a direct antiviral role of Hippo signaling on SARS-CoV-2 infection and disease pathogenesis processes, which can be therapeutically targeted. A schematic diagram of our working hypothetical model is shown in Figure 3G.

In conclusion, our results show that SARS-CoV-2 infection caused activation of the Hippo signaling pathway in pluripotent stem cell-derived cardiomyocytes (PSC-CMs), human primary lung air-liquid interface (ALI) cultures, and human airway epithelial cells (Calu-3). These *in vitro* cultures are efficiently infected by SARS-CoV-2, consequently causing activation of immune, and inflammatory responses, and altered Hippo signaling cascade. shRNA-mediated partial knockdown of LATS1 and pharmacological modulation of core upstream MST1/2 kinases resulted in enhanced SARS-CoV-2 replication. Loss of function mutations in these antiviral kinases could enhance susceptibility to SARS-CoV-2 infections in human. While Verteporfin, a pharmacological inhibitor targeting downstream transactivator, YAP, significantly reduced viral replication and production. Moving forward, additional mechanistic investigations are required to elucidate the underpinnings of the Hippo pathway role in antiviral responses to RNA viruses, and the preclinical animal safety and efficacy studies of Verteporfin therapy for COVID-19. Taken together, our results indicate a direct antiviral role for Hippo signaling in SARS-CoV-2 infection, thus creating potentially new avenues this pathway can be pharmacologically targeted to treat COVID-19.

## METHODS/MATERIALS

### Ethics Statement

For human tissue procurement, large airways and bronchial tissues were acquired after lung transplantations at the Ronald Reagan UCLA Medical Center from deidentified normal human donors following Institutional Review Board (IRB) exemption. Pluripotent stem cell related work received UCLA ESCRO-IRB approval. All the SARS-CoV-2 live virus experiments were performed at the UCLA BSL3 High containment facility.

### Cell lines, hPSC-CM cultures and ALI cultures

Vero E6 cells were obtained from ATCC [VERO C1008 (ATCC^®^ CRL-1586^™^)] or DSMZ (Braunschweig, Germany). Cells were cultured in EMEM growth media containing 10% fetal bovine serum (FBS) and penicillin (100 units/ml). Human lung adenocarcinoma epithelial cell line (Calu-3) was purchased from ATCC (ATCCHTB-55) and cultured in Dulbecco’s Modified Eagles Medium (DMEM), supplemented with 20 % fetal bovine serum (FBS), 1 % L-glutamine (L-glu) and 1 % penicillin/streptomycin (P/S). Cells were incubated at 37°C with 5% CO_2_. Human pluripotent stem cell-derived cardiomyocytes (hPSC-CM) were derived from hESC line H9 using the previously described method^51^. Temporarily, hPSCs were maintained in mTeSR1 (STEMCELL Technology). Shortly after, RPMI1640 was supplemented with B27 minus insulin (Invitrogen) and used as differentiation medium. From Day 0-1, 6 μM CHIR99021 (Selleckchem) was added into differentiation medium. Next, Day 3-5, 5 μM IWR1 (Sigma-Aldrich) was added to differentiation medium. After Day 7, RPMI 1640 plus B27 maintenance medium was added. From Day 10-11, RPMI 1640 without D-glucose supplemented with B27 was transiently used for metabolic purification of CMs^52^. After selection, hPSC-CMs were replated for viral infection. Air-liquid interface (ALI) cultures derived from primary human proximal airway basal stem cells (ABSCs) were used as described previously^39^. 24-well 6.5mm trans-wells with 0.4mm pore polyester membrane inserts were used for culturing ALI cells. 500 μl ALI media (PneumaCult™-ALI Medium, STEMCELL Technologies) was used in the basal chamber for ALI cultures and cells were cultured at 37°C with 5% CO_2_. Use of these established cell lines for the study was approved by the Institutional Biosafety Committee at UCLA.

### shRNA-mediated gene silencing

hPSC-CM cells (1 × 10^5^ cells/well) were added in a 48-well plate. The pLKO.1-puro sh-RNA targeting YAP1 (5’-CCGGCCCAGTTAAATGTTCACCAATCTCGAGATTGGTGAACATTTAACTGGGTTTTTG-3’), LATS1 (5’-CCGGCAAGTCAGAAATCCACCCAAACTCGAGTTTGGGTGGATTTCTGACTTGTTTTT-3’) or pLKO.1-puro Non-Targeting shRNA Control (Sigma-Aldrich) lentiviral particles were added to the cells. At 72 hours post-transduction, SARS-CoV-2 (Isolate USA-WA1/2020) with a multiplicity of infection (MOI) of 0.01, was added. At 48 hours post-infection, cells were fixed in 4% paraformaldehyde for IHC studies or cell protein lysates were collected for western blot analysis.

### Pharmacological modulation of Hippo signaling pathway in infected cells

hPSC-CM and Calu-3 cells were seeded on 48-well plates or 96-well plates, respectively. hPSC-CM cells were pretreated with 4-((5,10-dimethyl-6-oxo-6,10-dihydro-5H-pyrimido[5,4-b]thieno[3,2-e][1,4]diazepin-2-yl)amino)benzenesulfonamide (XMU-MP-1 at 10 μM) and Calu-3 cells were treated with XMU-MP-1 (10 μM) or Verteporfin (1 μM). Cells were pretreated with drugs for 24 hours, then infected with SARS-CoV-2 inoculum (MOI 0.1). DMSO vehicle treated cells, with or without viral infections, were included as controls. At 48 hpi, the cells were fixed with 4% PFA or lysed with RIPA buffer for proteins. The cell-free culture supernatants were collected from Calu-3 for viral titer measurement. Fixed cells were immunostained with anti-spike antibody (NR-616 Monoclonal Antibody) to assess virus replication.

### Virus

SARS-Related Coronavirus 2 (SARS-CoV-2), Isolate USA-WA1/2020, and Isolate hCoV-19/USA/MD-HP05647/2021 (Lineage B.1.617.2; Delta variant) were obtained from BEI Resources of National Institute of Allergy and Infectious Diseases (NIAID). All the studies involving live virus was conducted in UCLA BSL3 high-containment facility. SARS-CoV-2 was passaged once in Vero E6 cells and viral stocks were aliquoted and stored at −80°C. Virus titer was measured in Vero E6 cells by established plaque assay or TCID50 assay.

### SARS-CoV-2 Infection

Calu-3 cells were seeded at 30 × 10^3^ cells per well in 0.2 ml volumes using a 96-well plate and hPSC-CMs were replated at 1 × 10^5^ cells per well in a 48-well plate. Viral inoculum (MOI of 0.01 and 0.1; 100 μl/well) was added using serum free base media. The conditioned media from each well and condition was removed and 100 μl of prepared inoculum was added onto cells. After 1 hour incubation at 37°C with 5% CO2, inoculum was replaced for Calu-3 cells with serum supplemented media (200 μl per well) and for hPSC-CM, cell culture medium was replaced with RPMI 1640 + B27 supplement with insulin. For ALI cultures, 100 μl of viral inoculum prepared in PneumaCult media was added to the apical chamber of ALI culture insert and incubated for 1 hour at 37°C with 5% CO2. For mock infection, PneumaCult media (100 μl/well) alone was added. The inoculum was dispersed throughout the hour every 15 minutes by gently tilting the plate sideways. At the end of incubation, the inoculum was removed from the apical side for ALI cultures to maintain the air liquid interface^39^. At selected timepoints, cells were fixed with 4% PFA, collected by 1xRIPA for protein analysis, and/or supernatant collected for viral titer. Viral infection was examined by immunostaining or western blot analysis using SARS-CoV-2 antibodies [BEI Resources: NR-10361 polyclonal anti-SARS coronavirus (antiserum, Guinea Pig), and NR-616 monoclonal anti-SARS-CoV S protein (Similar to 240C) SARS coronavirus].

### Viral Titer by TCID50 *(Median Tissue Culture Infectious Dose)* Assay

Viral production by infected cells was measured by quantifying TCID50 as previously described^53^ methodology for quantifying TCID50.10 ×10^3^cells/well of Vero E6 cells were seeded in 96-well plates. The next day, culture media samples of supernatant collected from Calu-3 cells were at 48 hour time point, was subjected to 10-fold serial dilutions (10^1^ to 10^8^) and inoculated onto Vero E6 cells. The cells were incubated at 37°C with 5% CO2 for 3 to 4 days to evaluate for presence or absence of viral CPE. After measuring the percent infected dilutions immediately above and immediately below 50%, the TCID50 was calculated based on the method of Reed and Muench.

### Histopathology

COVID 19 patient autopsy lung samples were procured from the UCLA Translational Pathology Core Lab for research use. Samples were processed and sectioned, after a pathologist confirmed the section quality by H&E staining, and subsequent confirmation of COVID-19 positivity by RNAscope V-nCoV2019-S probe (ACD, Cat#: 848568, ready to use). Immunohistochemistry stainings were also performed on this lung tissue: Paraffin-embedded sections were cut at 4μm thickness and paraffin was removed with xylene and the sections were rehydrated through graded ethanol. Endogenous peroxidase activity was blocked with 3% hydrogen peroxide in methanol for 10 min. Heat-induced antigen retrieval (HIER) was carried out for all sections in AR9 buffer (AR9001KT Akoya) using a Biocare decloaker at 95°C for 25 min. The slides were then stained with YAP (S127) antibody (Cell Signaling, 13008, 1-100) and CD68 antibody (Dako, m0876, 1-200) at 4 degree overnight, the signal was detected using Bond Polymer Refine Detection Kit (Leica Microsystems, catalogue #DS9800) with a diaminobenzidine reaction to detect antibody labeling and hematoxylin counterstaining.

### Immunohistochemistry

Cells were fixed with 4%PFA for 30-60 minutes or methanol (incubated in −20°C freezer until washed with 1XPBS). Cells were washed 3 times with 1x PBS before permeabilizing with blocking buffer (0.3% Triton X-100, 2% BSA, 5% Goat Serum, 5% Donkey Serum in 1 X PBS) for 1 hour at room temperature. After adding specific primary antibodies (Refer to Supplementary Table 1), cells were incubated overnight at 4°C. The next day, cells were washed with 1X PBS three times and incubated with respective secondary antibody (Refer to Supplementary Table 1) at room temperature for 1 hour. DAPI (4’,6-Diamidino-2-Phenylindole, Dihydrochloride) (Life Technologies) was used to stain nuclei at a dilution of 1:5000 in 1 X PBS. Image acquisition was done using Leica DMi1 fluorescent microscopes and using the Leica Application Suite X (LAS X) The LSM 700 confocal microscopes and Zeiss Software programs available at the UCLA Eli & Edythe Broad Center of Regenerative Medicine & Stem Cell Research Microscopy Core at Center for Health Sciences Building was used for capturing images as well.

### Western Blot analysis

Cells were lysed in 1x RIPA 50 mM Tris pH 7.4, 1% NP-40, 0.25% sodium deoxycholate, 1 mM EDTA, 150 mM NaCl, 1 mM Na3VO4, 20 Mm or NaF, 1mM PMSF, 2 mg ml^-1^ aprotinin, 2 mg ml^-1^ leupeptin and 0.7 mg ml^-1^ pepstatin or Laemmli Sample Buffer (Bio Rad, Hercules, CA). Cell lysates were resolved by SDS-PAGE using 10% gradient gels and transferred to a 0.2 μm PVDF membrane. Subsequently, the membranes were blocked with 5% skim milk and 0.1% Tween-20 in 1x TBST (0.1% Tween-20) at room temperature for 1 hour. The membranes were then probed with respective monoclonal antibodies and detected by Thermo Scientific SuperSignal West Femto Maximum Sensitivity Substrate.

### Image Analysis/Quantification

The confocal images were obtained using the Leica Application Suite X (LAS X) and/or the Zeiss LSM 700 Confocal Microscopy by Zeiss Software Program with maximum intensity projection feature. Using a double blinded approach to count the positively stained cells, Image J’s plugin Cell Counter program was used. 3-4 independent images per condition of the hPSC-CM were analyzed. For all the analysis, we used confocal images acquired at 63X objective and fluorescent microscope images acquired at 20 and 40x. Total number of cells in each image was obtained by manually counting all the DAPI stained nuclei using ImageJ program (Version 1.8.0; https://imagej.nih.gov/ij/index.html). Total cell count was then used for normalization and to calculate the percentage of individual marker positive cell populations in respective images. The mean percentage of positively stained cells from 3-8 independent images (about 150-1000 cells total) were quantified and presented as graph format.

### RNA sequencing data analysis

High throughput sequencing dataset of five healthy and autopsy lung samples from COVID-19 patients (L2, L3 and L4 from case 6; L1 from case 7 and L1 from case 10) deceased due to SARS-CoV-2 infection were obtained from the NCBI Gene Expression Omnibus (GEO) under the accession number GSE150316^54^. The SARS-CoV-2 infected pluripotent stem cell-derived cardiomyocytes gene expression data used in this study were retrieved from GEO under the accession number GSE150392^46^. The raw read counts per gene were used as inputs for differential expression gene analysis using DESeq2 v1.28.1 in R v4.0.3^55^. Median of ratios method was used to normalize expression counts for each gene in all experimental samples. Each gene in the samples was fitted into a negative binomial generalized linear model. Genes were considered as differentially expressed (DEG) if they were supported by a false discovery rate (FDR) p<0.01. The heatmaps were prepared using shinyheatmap web interface^56^. RNA-seq data generated from hiPSC-CM used in this study was retrieved at Gene Expression Omnibus with accession number GSE150392^57^.

High throughput sequencing dataset of five healthy and autopsy lung samples from three patients (L2, L3 and L4 from case 6; L1 from case 7 and L1 from case 10) deceased due to SARS-CoV-2 infection were obtained from GEO under the accession number GSE150316 (https://www.ncbi.nlm.nih.gov/geo/query/acc.cgi?acc=GSE150316). The raw read counts per gene were used for differential expression gene analysis using v1.28.1 in R v4.0.3. DEG were considered if they supported by a FDR p<0.01. The Kyoto Encyclopedia of Genes and Genomes (KEGG) pathway database was used to examine the hippo signaling pathway in the DEG datasets^58^.

## Data analysis

All statistical testing was performed at the two-sided alpha level of 0.05. To test statistical significance, unpaired student’s *t*-test was used to compare two groups (uninfected vs. infected). GraphPad Prism software, version 8.1.2 (GraphPad Software, US) was used.

## Acknowledgments

We are grateful to Barbara Dillon, UCLA High Containment Program Director for BSL3 work. We thank Nate Price for assistance with manuscript editing and proofreading, and Nikhil Chakravarty for design and production of illustrations. This study is supported by University of California, Los Angeles (UCLA) David Geffen School of Medicine (DGSOM), and Broad Stem Cell Research Center (OCRC #20-15), UCLA W.M. Keck Foundation COVID-19 Research Award, and National Institute of Health awards 1R01EY032149-01 to V.A.,1R01DK132735-01 to V.A. and A.D., and 1R01CA208303 to B.G. The research of K.M. is supported by the National Institute of Health grants R01AI145044 and U19AI149504. A.R. is supported by the Tata Institute for Genetics and Society. The following reagents were obtained through BEI Resources, NIAID, NIH: Monoclonal Anti-SARS-CoV S Protein (Similar to 240C), NR-616; Polyclonal Anti-SARS Coronavirus (antiserum, Guinea Pig), NR-10361; SARS-Related Coronavirus 2, Isolate USA-WA1/2020, (NR-52281); SARS-CoV-2, Isolate hCoV-19/USA/PHC658/2021 (Delta Variant) (NR-55611).

## Author contributions

Garcia Jr. G: Conception and design, Collection and/or assembly of data. Data analysis and interpretation, and Manuscript writing.

Wang Y., Ignatius Irudayam J., Jeyachandran AV., Cario SC., Sen C., Li S.: Conducted experiments, Data analysis and interpretation.

Kumar A, Nielsen-Saines K., French SW., Shah P., Morizono K., Gomperts B., Deb A., Ramaiah A.: Experimental design, Data analysis, interpretation and Manuscript writing.

Arumugaswami V: Conception and design, Data analysis and interpretation, Manuscript writing and Final approval of manuscript.

## Data and materials availability

All relevant data regarding this manuscript is available from the above listed authors. Supplementary information is available for this paper. The gene expression data used in this study were retrieved at Gene Expression Omnibus with accession numbers GSE150316 ^54^ and GSE150392^57^. Correspondence and requests for materials should be addressed to Vaithilingaraja Arumugaswami.

## SUPPLEMENTARY FIGURES

**Supplementary Figure 1.**
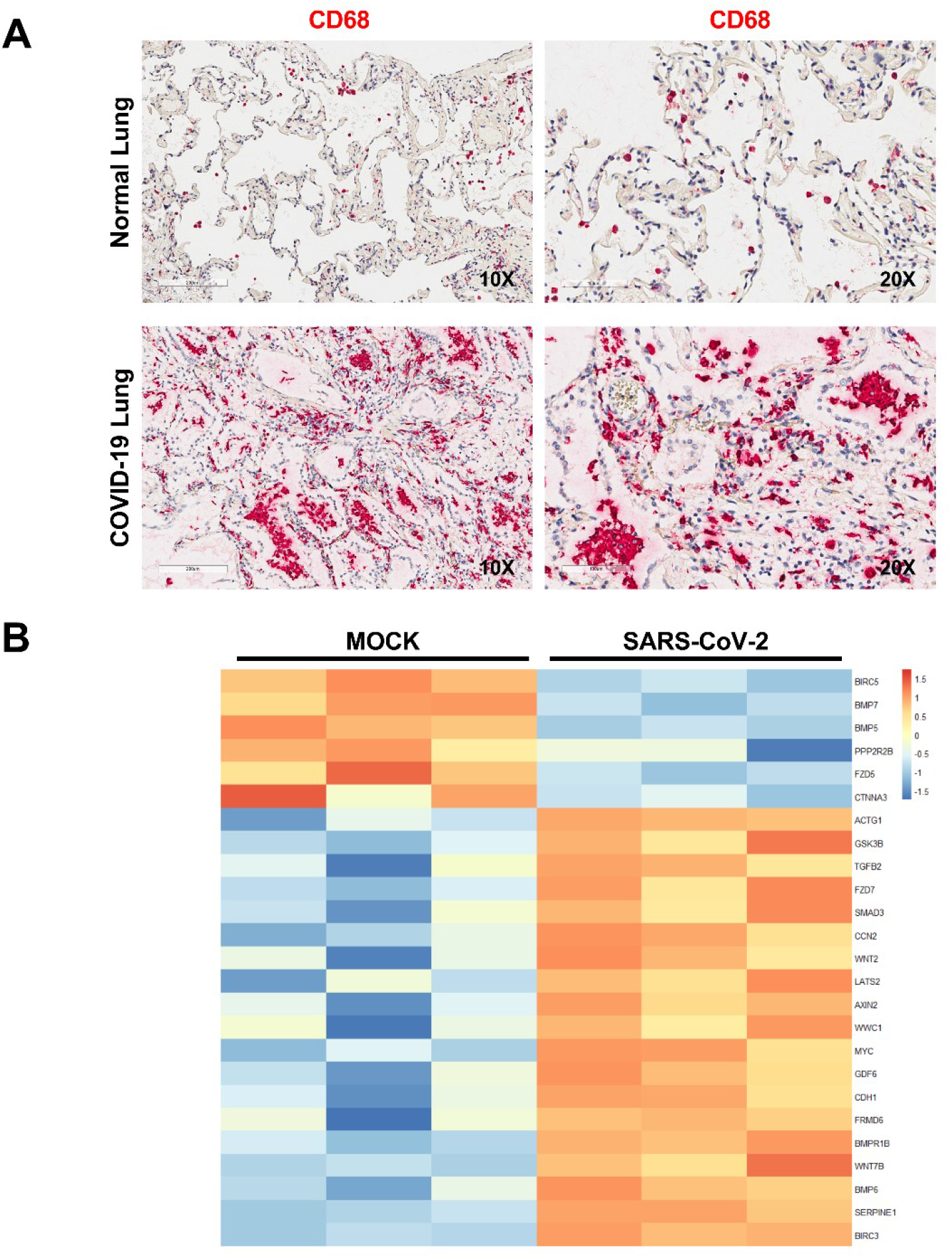
(A) Immunohistochemistry of COVID-19 lung autopsy tissue shows high level of CD68 positive inflammatory cells (red). Images are obtained at 10x and 20x magnifications. (B) Transcriptome analysis of control and SARS-CoV-2 infected human induced PSC-CMs at 3 dpi. Heatmap depicting *Z* scores as expression levels of the 25 differentially expressed genes (*p* <0.01) involved in Hippo signaling pathway. Blue and red colors represent downregulated and upregulated genes, respectively. The gene expression data was retrieved at Gene Expression Omnibus with accession number GSE150392.

**Supplementary Figure 2.**
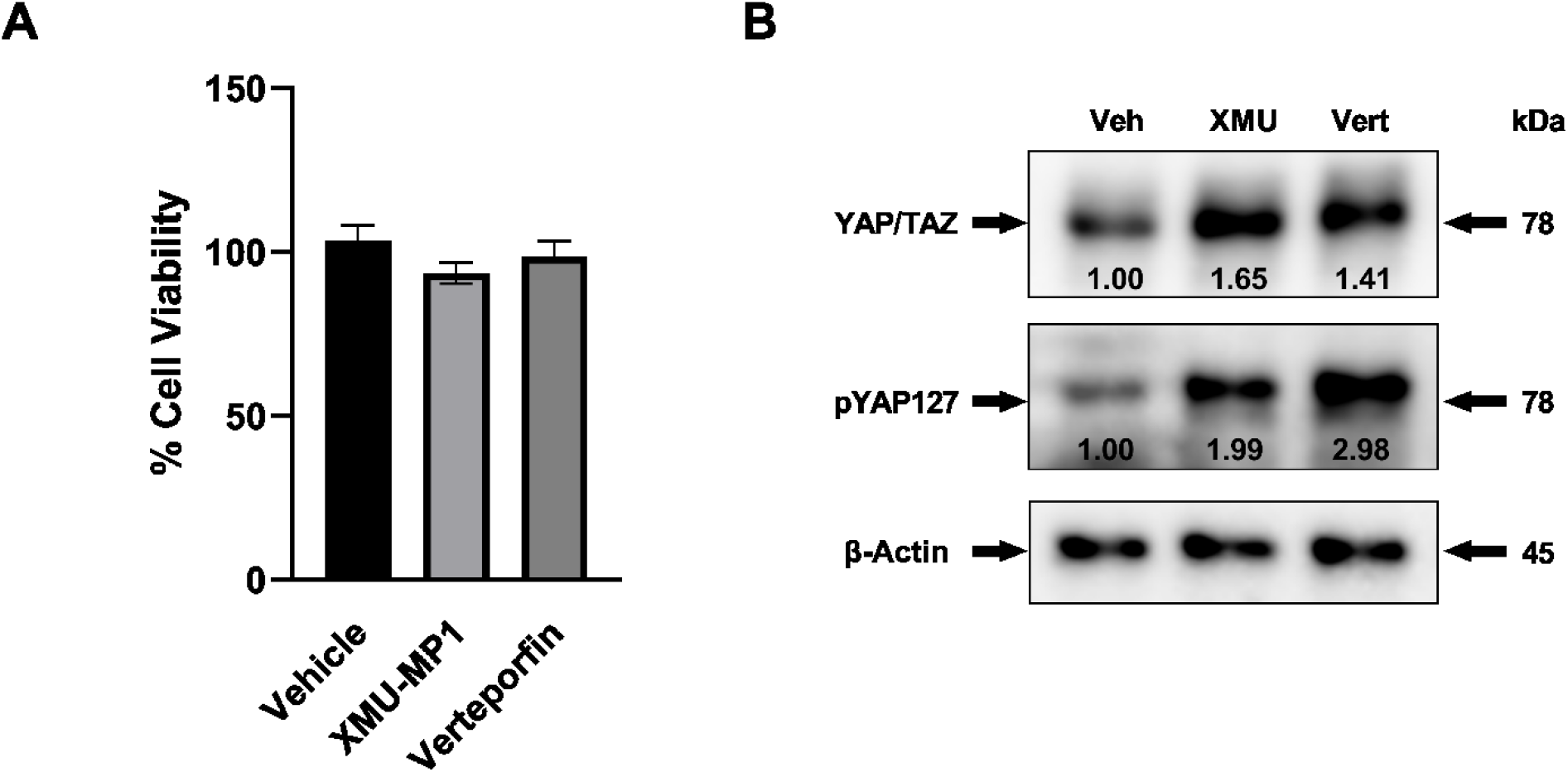
(A) Graph shows the percent cytotoxicity of Calu-3 cells 72 hours post-treatment with DMSO (Vehicle), XMU-MP1 (10μM), and Verteporfin (1 μM). CellTiter-Glo Luminescent Cell Viability Assay was performed as per the manufacturer (Promega, USA) recommendation. (B) Western blot analysis shows total and phosphorylated YAP at 72 hours-post treatment with indirect and direct acting inhibitors, XMU-MP-1 and Verteporfin, respectively. Note: XMU-MP-1 treatment enhances YAP/TAZ level compared to vehicle, whereas Verteporfin increases phosphorylated YAP (S127) levels. Representative data from two independent experiments is shown.

## SUPPLEMENTAL TABLES

**Supplementary Table 1:** Reagents or resources used in this study.

| REAGENT/RESOURCE | SOURCE | IDENTIFIER |
| --- | --- | --- |
| **Antibodies** |  |  |
| Monoclonal anti-SARS-CoV S protein (Similar to 240C) antibody | BEI Resources Repository | Cat#NR-616 |
| Polyclonal anti-SARS coronavirus (antiserum) | BEI Resources Repository | Cat#NR-10361 |
| TBK1/NAK (D1B4) | Cell Signaling Technology | Cat#3504S |
| Phospho-TBK1/NAK (Ser172) | Cell Signaling Technology | Cat#5483S |
| YAP/TAZ (D24E4) | Cell Signaling Technology | Cat# 8418S |
| Phospho-YAP (Ser127) | Cell Signaling Technology | Cat#13008S |
| COVID V-nCoV2019-S probe | ACD | Cat#: 848568 |
| AC TUBULIN | Cell Signaling Technology | Cat#mAb#5335 |
| MUC5AC Mouse | Invitrogen | Cat#MA512178 |
| Anti-Cardiac Troponin T antibody | Abcam | Cat#ab45932 |
| Cleaved Caspase 3 | Cell Signaling Technology | Cat# 9661S |
| Cleaved caspase-3 rabbit monoclonal antibody, clone D175 | Cell Signaling | Cat#9661S |
| Goat anti-Mouse IgG (H+L) Cross-Adsorbed Secondary Antibody, Alexa Fluor 555 | Thermo Fisher Scientific | Cat#A21422 |
| IgG (H+L) Cross-Adsorbed Goat anti-Rabbit, Alexa Fluor 488, Invitrogen | Thermo Fisher Scientific | Cat#A11008 |
| IgG (H+L) Cross-Adsorbed Goat anti-Human, Alexa Fluor 488, Invitrogen | Thermo Fisher Scientific | Cat#A11013 |
| Goat anti-Guinea Pig IgG (H+L) Highly Cross-Adsorbed Secondary Antibody, Alexa Fluor 488 | Thermo Fisher Scientific | Cat#A11073 |
| Monoclonal Anti-Beta-Actin, Clone AC-74 produced in mouse | MilliporeSigma | Cat#A2228 |
| Phospho-Stat1 (Tyr701) (58D6) Rabbit mAb | Cell Signaling | Cat#9167S |
| Stat1 (D1K9Y) Rabbit mAb | Cell Signaling | Cat#14994 |
| CD68 | Dako | Cat#m0876 |
| **Bacterial and Virus Strains** |  |  |
| SARS-Related Coronavirus 2 (SARS-CoV-2), Isolate USA-WA1/2020 | BEI Resources Repository | Cat#NR-52281 |
| SARS-Related Coronavirus 2, Isolate hCoV-19/USA/MD-HP05647/2021 (Lineage B.1.617.2; Delta variant), | BEI Resources Repository | Cat#NR-55672 |
| SHCLNG MISSION shRNA YAP1 Bacterial Clone | Sigma-Aldrich | Cat#<br>TRCN0000107267 |
| SHCLNG MISSION shRNA LATS1 Bacterial Clone | Sigma-Aldrich | Cat#<br>TRCN0000001779 |
| **Chemicals, Peptides, and Recombinant Proteins** |  |  |
| Regular Fetal Bovine Serum | Corning | Cat#35010CV |
| Eagle's Minimum Essential Medium (MEM) | Corning | Cat#10009CV |
| Penicillin-Streptomycin (10,000 U/mL) | Gibco | Cat#15140122 |
| L-Glutamine (200 mM) | Gibco | Cat#25030081 |
| Puromycin Dihydrochloride | Gibco | Cat#A1113803 |
| PneumaCult™-ALI Medium | STEMCELL Technologies | Cat#05021 |
| AR9 BUFFER, 10X | Akoya Biosciences | Cat#AR9001KT |
| Dimethyl sulfoxide | MilliporeSigma | Cat#D2650 |
| RPMI 1640 | Thermo Fisher Scientific | Cat#11875093 |
| B27 supplement with insulin | Thermo Fisher Scientific | Cat#17504044 |
| Methanol (Histological) | Thermo Fisher Scientific | Cat#A433P4 |
| 16% Paraformaldehyde (formaldehyde) aqueous solution | Electron Microscopy Sciences | Cat#15710 |
| Dulbecco's Phosphate-Buffered Salt Solution 1X | Corning | Cat#21030CV |
| Perm/Wash Buffer | BD Biosciences | Cat#554723 |
| DAPI (4',6-Diamidino-2-Phenylindole, Dihydrochloride) | Thermo Fisher Scientific | Cat#D1306 |
| SuperBlock™ (PBS) Blocking Buffer | Thermo Fisher Scientific | Cat#37515 |
| Bovine Serum Albumin | MilliporeSigma | Cat#A9418 |
| Normal Donkey Serum | Jackson ImmunoResearch | Cat#017-000-121 |
| Normal Goat Serum | Cell Signaling | Cat#5425S |
| Triton-X 100 | MilliporeSigma | Cat#T9284 |
| XMU-MP-1 | MilliporeSigma | Cat# SML2233-5MG |
| Verteporfin | MilliporeSigma | Cat#SML0534-5MG |
| **Commercial Assays** |  |  |
| Bond Polymer Refine Detection Kit | Leica Microsystems | Cat#DS9800 |
| CellTiter 96® Non-Radioactive Cell Proliferation Assay (MTT) | Promega | Cat#G4000 |
| **Experimental Models: Cell Lines** |  |  |
| VERO C1008 [Vero 76, clone E6, Vero E6] | ATCC | Cat#CRL-158 |
| Calu-3 | ATCC | Cat#HTB-55 |
| hPSC derived cardiomyocyte | University of California, Los Angeles (Li et al., 2021) | N/A |
| Normal human bronchial epithelial cells | Lonza | N/A |
| **Deposited Data** |  |  |
| RNA-Seq of Human iPSC-cardiomyocytes infected with SARS-CoV-2 | Gene Expression Omnibus | Accession Number: GSE150392 |
| **Software and Algorithms** |  |  |
| GraphPad Prism 8 | GraphPad | N/A |
| Multi-Point Tool (Cell Counter) | ImageJ | N/A |

